# MicroRNA regulation of BMP signaling; cross-talk between endothelium and vascular smooth muscle cells

**DOI:** 10.1101/395541

**Authors:** Charlene Watterston, Lei Zeng, Abidemi Onabadejo, Sarah J Childs

**Affiliations:** Alberta Children’s Hospital Research Institute, and Department of Biochemistry and Molecular Biology, Cumming School of Medicine, University of Calgary, Calgary AB, Canada, T2N 4N1.

## Abstract

Vascular smooth muscle cells (vSMC) are essential to the integrity of blood vessels, and therefore an attractive target of therapeutics aimed at improving vascular function. Smooth muscle cells are one of the few cell types that maintain plasticity and can switch phenotypes from differentiated (contractile) to de-differentiated (synthetic) and vice versa. As small regulatory transcripts, miRNAs act as genetic ‘fine tuners’ of posttranscriptional events and can act as genetic switches promoting phenotypic switching. The microRNA *miR26a* targets the BMP signalling effector, *smad1*. We show that loss of *miR26* leads to hemorrhage (a loss of vascular stability) *in vivo*, suggesting altered vascular differentiation. Reduction in *miR26a* levels increases *smad1* mRNA and phospho-Smad1 (pSmad1) levels. We show that increasing BMP signalling by overexpression of *smad1* also leads to hemorrhage and that normalization of Smad1 levels through double knockdown of *miR26* and *smad1* rescues hemorrhage suggesting a direct relationship between *miR26* and *smad1* and vascular stability. Using a BMP genetic reporter and pSmad1 staining we show that the effect of *miR26* on vascular instability is non-autonomous; BMP signalling is active in embryonic endothelial cells, but not in smooth muscle cells. Nonetheless, increased BMP signalling due to loss of *miR26* results in an increase in *acta2*-expressing smooth muscle cell numbers and promotes a differentiated smooth muscle morphology. Taken together our data suggests that *miR26* modulates BMP signalling in endothelial cells and indirectly promotes a differentiated smooth muscle phenotype. Our data also suggests that crosstalk from BMP-responsive endothelium to smooth muscle is important for its differentiation.

## Introduction

Vascular smooth muscle cells (VSMCs) provide structural integrity to the vessel wall. Guided control of signalling cascades, including P*latelet Derived Growth Factor (PDGF), Notch, and Transforming Growth Factor-β/Bone morphogenic Protein (TGF-β/BMP)*, recruit and induce a perivascular niche of cells termed mural cells (comprising of pericytes and immature vSMCs) to create a two-layered vessel wall with an internal endothelial cell lining and a muscle cell covering (Gaengel et al. 2009; Lamont et al. 2010; Liu et al. 2007). Once the vSMCs surround the vessel they begin depositing extracellular matrix proteins such as *laminin, collagen IV* and *fibulins* to support cellular contacts (Rensen et al. 2007). vSMCs then take on a mature contractile phenotype that stabilizes the underlying endothelial cells through induction of quiescence, expression of junctional and attachment protein, and expression of contractile proteins to provide myogenic tone (Lamont et al. 2010; David et al. 2008; Owens et al. 1996; Whitesell et al. 2014).

vSMCs maintain phenotypic plasticity and can undergo a phenotypic switches from a quiescent contractile state to a proliferative synthetic state in response to various cellular stimuli (Rensen et al. 2007; Hendrix et al. 2005). Contractile vSMC are defined by an elongated, spindle-shaped morphology and low rates of proliferation. The expression of key differentiation markers such as smooth muscle (α)-actin (Acta2), smooth muscle β-myosin heavy chain (Myh11), and transgelin (Sm22α) allows vSMCs to perform their contractile function and provide vascular tone. In contrast, rhomboid, immature (synthetic) vSMCs are highly proliferative, produce ECM proteins and have reduced expression of contractile genes (Owens et al. 2004; Mack & Owens 1999; Miano et al. 1994; Lehoux et al. 2006).

Cumulative studies have demonstrated that BMPs and Smad1 signaling have roles in modulating vSMC plasticity (reviewed by Cai et al. 2012). Aberrant vSMCs phenotype switching plays a critical role in the pathogenesis of vascular diseases, such as hereditary hemorrhagic telangiectasia (HHT) and pulmonary arterial hypertension (PAH), with defective BMP signalling. The diseases affect both endothelial function and vSMC differentiation (Li et al. 1996; El-Bizri et al. 2008; Torihashi et al. 2009; Benjamin et al. 1998; Lan et al. 2007; Roman et al. 2002; Beppu et al. 2000; Mishina et al. 1995). Murine homozygous null mutants for BMPR-1a (Activin like kinas 3, ALK3) or the type II receptor BMPR-2 (which is mutated in human patients with PAH) (Paul et al. 2005; Orvis et al. 2008) and their ligand Bmp4 or downstream co-SMAD4 are embryonic lethal, and present with vascular deformities attributable to a loss of Smad1 mediated signalling (Sirard et al. 1998). Mutations in ALK1 led to HHT2, a disease characterized by arteriovenous malformations (AVMs) (McDonald et al. 2011). Deletion of Alk-1 in mice leads to cranial hemorrhages, AVM-like fusion of micro-vessel plexi, dilation of large vessels and reduced coverage of vessels by vSMCs (Park et al. 2006). In fish, while disruption of *Alk1* signalling results in pathological arterial enlargement and maladaptive responses generate AVMs, additional vSMCs defects have not been assessed (Corti et al. 2011).

As small noncoding RNAs, microRNAs (miRs) are genetic cues that regulate gene expression of key vSMC marker genes to control vSMC dynamics. (Albinsson et al. 2010; Ji et al. 2007; Xie et al. 2011; Zhang 2010). A number of miRNAs have been identified as modulators of the vSMC phenotype *in vitro* and *in vivo*, including *miR-145, miR-21, miR-221, miR-222* and *miR-146a* (Zeng et al. 2009; Rangrez et al. 2011; Xu et al. 2011; Sarkar et al. 2010; Ali et al. 2015; Sun et al. 2011; Wang et al. 2010; Xin et al. 2009; Cordes et al. 2009). (Xin et al. 2009; Cordes et al. 2009; Zeng et al. 2009; Wang et al. 2010; Sarkar et al. 2010; Rangrez et al. 2011; Bai et al. 2011; Sun et al. 2011; Xu et al. 2011; Ali et al. 2015) We previously showed that miR-145 promotes visceral smooth muscle differentiation via controlling cross-talk between visceral epithelial cells and smooth muscle (Zeng et al. 2009; Zeng & Childs 2012).

In this study we investigated the role of *microRNA26a* (*miR26a*) in regulating vSMC dynamics the zebrafish model of vessel stabilization. *miR26* has been identified as a potential regulator of vSMC proliferation, migration and differentiation of vSMCs *in vitro* (Icli et al. 2014; Leeper et al. 2011; Bai et al. 2011; Yang et al. 2017). miR-26a expression is altered during abdominal aortic aneurysm (AAA) and neointimal lesion formation (Leeper et al. 2011; Yang et al. 2017), and although primarily categorized as endothelial defects, the pathological progression and compromised vessel structure is attributed to a shift in vSMC phenotypes. However, the significance and cell autonomy of these interactions in an intact animal and in context of developing vSMC are largely unknown. Using a combination of transient knockdown methods to understand the effect of decreasing *miR26a* levels in vivo, we show that *miR26a* acts within a BMP responsive pathway to fine tune vascular smooth muscle cell maturation via targeting *smad1*. However, active BMP signalling was observed within endothelium in vivo, and not in smooth muscle cells. Together the evidence suggests that *miR26a* has a role in regulating blood vessel stabilization via a non-autonomous mechanism.

## Materials and Methods

### Zebrafish

Zebrafish (*Danio Rerio)* embryos were collected and incubated at 28.5 °C in E3 embryo medium and staged in hours post-fertilization (hpf) or days post fertilization (dpf). Endogenous pigmentation was inhibited from 24 hpf by the addition of 0.003% 1-phenyl-2-thiourea (PTU, Sigma-Aldrich) in E3 embryo medium. All animal procedures were approved by the University of Calgary Animal Care Committee. The fluorescent transgenic endothelial mCherry-expressing *Tg(kdrl:mcherry)*^*ci*5^, GFP-expressing *Tg(kdrl:EGFP)*^*la116*^and *Tg(fli:nEGFP)*^*y7*^(Roman et al. 2002) report endothelial expression, *Tg(acta2:GFP)*^*ca7*^ and *Tg(acta2:mCherry)*^*ca8*^ report smooth muscle expression (Whitesell et al. 2014) and BMP-reporter fish *Tg(BRE-AAVmlp:eGFP)*^*mw29*^ [hereafter BRE:eGFP] report active BMP signaling (Collery & Link 2011).

### Morpholino knockdown, CRISPRi and mRNA overexpression

Both MO and mimic were injected into one-to four-cell stage embryos within recommended dosage guidelines (Bill et al. 2009; Schulte-Merker & Stainier 2014). Injected doses were 1ng/embryo for *miR26* MO, Scr. control *miR26*, and *smad1* MO. Morpholinos (MO) were obtained from Gene Tools LLC Corvalis, OR, USA. *mir-26a* MO blocks the mature microRNA (5-AGCCTATCC*TG*GATTACT*TG*AAC-3’), *miR26a* Scrambled control has 6bp mismatch (5’-ACCGTATCG*TG*CATTACTTCAAC-3’), and *smad1* MO blocks Smad1 translation (5’-AGGAAAAGAG*TG*AGG*TG*ACATTCAT-3’). For rescue experiments embryos were first injected with *miR26*MO and then *smad1* MO. To control for non-specific neural cell death that occurs from nonspecific activation of p53 with morpholinos, a standard p53 MO was co-injected with high dose morpholino to establish dosage curve. Hsa *miR26a* miRIDIAN mimic was obtained from Dharmacon and injected in a dose of 3ng/embryo.

For CRISPRi mediated knockdown of *miR26a*, sgRNA target were designed using CHOPCHOP (Montague et al., 2014) to target the seed sequence of *miR26a* family members to reduce *miR26* processing (*miR26a*-1, 26a-2, 26a-3, are independent genes located on different chromosomes. *miR26b* differs by one nucleotide). To generate sgRNA we followed a method established by (Narayanan et al. 2016). 10 μmol of forward primer and 50 μmol of a universal reverse primer (IDT Oligos, Yale) were annealed and filled in (Montague et al. 2014), purified (Qiagen PCR purification kit) and in vitro transcribed (T7 mMessage machine, Ambion). Primer sequence information is in supplemental table 1. Zebrafish codon optimized dCas9 plasmid (Rossi et al. 2015) was linearized with XbaI and in vitro transcribed using Ambion Maxi Kit (Life Technologies Inc., Burlington, ON), and RNA purified using an RNeasy Mini Kit (Qiagen). Zebrafish embryos at the one-cell stage were injected with 200pg of a solution containing 75 ng/μ l of sgRNA with 150 ng/μl of Cas9 mRNA.

For overexpression of *smad1*, mRNA was in vitro transcribed as described (McReynolds et al. 2007; gift from Todd Evans Lab) using mMessage mMachine (Life Technologies Inc., Burlington, ON). 40pg of mRNA was injected per embryo at the 1 cell stage.

### Small molecule inhibition

K02288 was used at a dose of 15µM (SML1307, Sigma). DSMO (D8418, Sigma) was used as a vehicle and control. Drug stocks were heated for 20min at 65C and then diluted in E3 embryo medium. Drug or control was applied to the from 52hpf until 4dpf. Embryos were grown at 28.5C in the dark until imaging, and drug changed once.

### qPCR analysis

Total RNA from 5-10 zebrafish embryos (or heads) per treatment group was isolated using the RNeasy Mini kit for mRNA or the miRNAs Mini kit for miRNA (Qiagen). For microRNA expression, 100ng of total RNA from each sample were reverse transcribed using the miRCURY LNA™ Universal RT cDNA Synthesis Kit and expression assayed using the miRCURY LNA™ Universal RT microRNA PCR System (Qiagen). The ΔΔCt method was used to calculate the normalized relative expression level of a target gene from triplicate measurements. Experiments were repeated independently at least three times. Primer sequences are listed in Table S1, expression levels normalized to that of miR-103a-3p or miR122 for miRNA expression.

### In situ hybridization and Immunostaining

Embryos were fixed in 4% paraformaldehyde in PBS with 0.1% Tween-20 at 4 °C overnight, followed by 100% methanol at −20 °C. For wholemount in situ hybridization embryos staining Digoxigenin (DIG)-labeled antisense RNA probes. Probes for *smad1* (construct describe by McReynolds et al. 2007) *sm22a, acta2, myh11a* were synthesized from PCR fragments amplified from embryonic zebrafish cDNA using the primers listed in supplementary table 1. Probes were synthesized by using SP6 or T7 RNA polymerase (Roche). *miR26a* double-DIG-labeled LNA probe was obtained from Exiqon.

Phospho-SMAD1/5/9 (pSMAD1/5/9) was detected with Rabbit anti-Phospho-Smad1 (Ser463/465)/Smad5(Ser463/465)/Smad9(Ser426/428) (1:400; Cell Signaling Technology) using an antigen retrieval protocol, GFP was detected with mouse anti-GFP antibody, JL8 (1:500, Clontech) and detected with Alexafluor 647 or 488 secondary antibodies (1:500; Invitrogen Molecular Probes).

### Imaging and statistical analysis

For imaging, embryos were immobilized in 0.004% (g/mol) Tricaine (Sigma) and mounted in 0.8% low melt agarose on glass bottom dishes (MatTek, Ashland, MA). Confocal images were collected on a Zeiss LSM 700 inverted microscope. Image stacks were processed in Zen Blue and are presented as maximal intensity projections. For cell counts images were converted to 16-bit images using ImageJ and threshold adjusted to allow counting of cells over a region of the VA from the anterior bulbous arteriosus to the most anterior PAA.

Data are expressed as mean ±SD. Two treatment groups were compared using Student’s t-test. Three or more treatment groups were compared by one-way ANOVA followed by post hoc analysis adjusted with a least significant-difference correction for multiple comparisons using GraphPad Prism version 7.00(La Jolla California USA).

## Results

### *miR26a* is expressed in vicinity of developing blood vessels

*miR26a* targets *smad1* and thereby directly regulates BMP signalling (model; Figure 1A). To observe the spatial gene expression pattern of *miR26a* in developing embryos we used in situ hybridization. At 48hpf, *miR26a* has a ubiquitous expression pattern (Figure 1 B-B’), however by 4 dpf the expression pattern becomes more localized in the ventral head of the embryo, with strong expression in the pharyngeal region, bulbous arteriosus and ventral aorta (Figure 1C). We observe that *miR26a* is expressed in and around the blood vessel endothelium (compare to kdrl:GFP transgenic signal; Figure 1C’, inset C’’) and may therefore play a role modulating BMP signalling in blood vessels.

**Figure 1:**
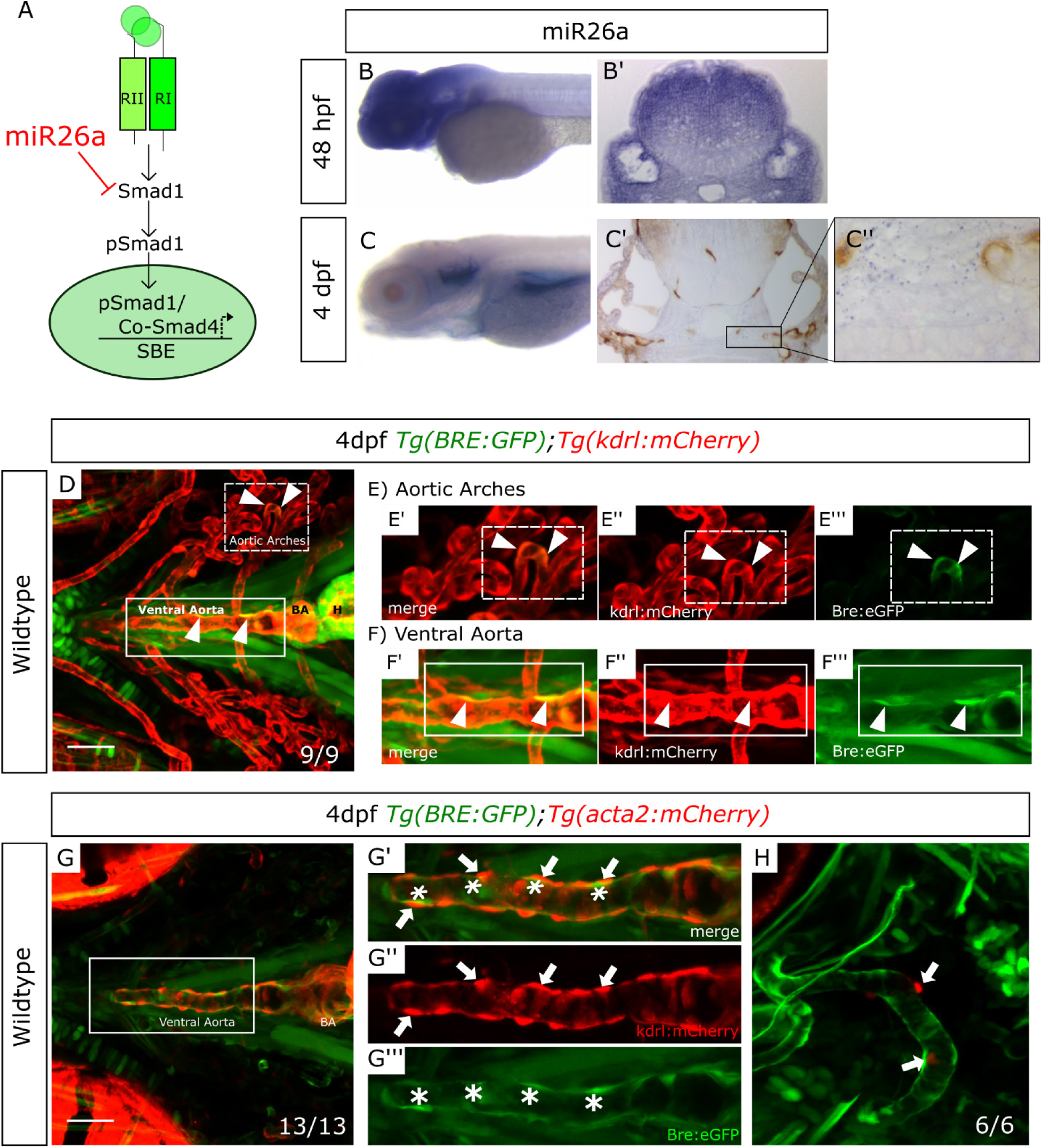
miR26 is expressed in the vicinity blood vessels; only endothelial cells have active BMP signaling. A) Model of how *miR26a* controls BMP signaling via direct targeting of *smad1*. In canonical Smad-mediated BMP signaling *Smad1* is phosphorylated by the serine-threonine kinase activity of type 1 BMP receptor, allowing it to associate and dimerize with the co-mediator Smad4 and translocate to the nucleus to control gene transcription. B) Lateral view of whole-mount in situ expression of *miR26a* at 2dpf shows ubiquitous expression pattern, with strong expression in the ventral head of the embryo. B’ Cross section of the head at 48 hpf. C) At 4dpf there is strong expression of miR26 in the pharyngeal arches, bulbous arterious and ventral aorta. C’) Cross section of the head showing miR26 expression in blood vessels (purple; punctate stain) compared with endothelial stain (brown; kdrl:GFP endothelial cell transgenic fish). C’’ enlargement of image in C’. D) Ventral view of the pharyngeal region of a 4 dpf double transgenic *Tg(BRE:eGFP);Tg(kdrl:mCherry)* embryo (BRE is in green; endothelial cells in red) shows BRE:eGFP expression within endothelial cells (arrowheads) on the E’-E’’’) aortic arches and F’-F’’’) ventral aorta. G and H) Ventral and lateral views of a 4 dpf double transgenic *Tg(BRE:eGFP); Tg(acta2:mCherry)* zebrafish shows that *acta2* positive cells are in direct contact with BMP-responsive endothelial cells but do not express BRE:eGFP. Scale bar= 50µm.

### BMP signalling is highly active in aorta endothelium of wild type embryos

We next tested where BMP signalling was active using an in vivo reporter of Smad1/5 activity. *Tg(BRE:eGFP]* transgenic fish encode eGFP driven by an upstream *Bmp Response Element (BRE)* that contains multiple short Smad-binding sites from the *id1* promoter, a major transcriptional target activated by canonical Bmp/Smad1 signaling (Collery & Link 2011; Laux et al. 2011). We crossed *Tg(BRE:eGFP]* to *Tg*(*kdrl*:mcherry) or *Tg(acta2:mCherry)* to localize active BMP signalling in endothelial and vSMCs respectively (Figure 1 D and G). vSMC cells first differentiate and begin to express the mature marker *acta2* between 3 and 4dpf (Whitesell et al. 2014; Georgijevic et al. 2007). Surprisingly, although *miR26a* has been implicated in controlling smad1 regulated vSMCs dynamics, transgenic *BRE:eGFP* signals were restricted to the endothelium of the vessel wall and co-expressed with *kdrl:mCherry* (Figure 1 D, E-D’ and F-F’). Differentiated *acta2:mCherry*-positive vSMCs lie directly adjacent to *BRE:eGFP*-expressing cells, with no detectable expression of *BRE:eGFP* in vSMCs on the ventral aorta or pharyngeal arch arteries (Figure 1 G-G’’). Similarly, *acta2:mCherry*-positive cells are in close association to pSmad1-positive endothelial cells but do not show pSmad1 staining (Supplemental Figure 1 B-B’). Together, our data suggests, that in early development, the endothelium, but not vSMCs respond to BMP signalling as visualized by two methods of detection.

### Knockdown of *mir26a* leads to upregulation of *smad1* and loss of vascular stability

The highly conserved miR-26 family constitutes miR-26a-1, miR-26a-2, and miR-26b (Bai et al. 2011) as identified by their seed sequences and accessory sequence. In zebrafish, and humans, *miR-26a-1, miR-26a-2* and *miR26a-3* have the same mature sequence, and only differ from the mature miR-26b sequence by two nucleotides (Icli et al. 2014; Griffiths-Jones et al. 2008). To investigate the role of *miR26a* in vascular development in vivo, we knocked down *miR26a* using an antisense morpholino that targeted the mature miRNA seed sequence of all three *miR26a* isoforms. A 6bp mismatch morpholino was used as a control. 1ng doses of morpholino were used, as suggested by current guidelines (Bill et al. 2009; Bedell et al. 2011). In parallel, we designed a second genetic knockdown approach using CRISPR interference (CRISPRi) (Long et al. 2015), to target the pri-miRNA hairpin structure using the complementary sequence to the mature miRNA (Figure 2 A). RT-qPCR shows a 22% reduction in miR26 following *miR26a* MO knockdown and 24% reduction with *miR26a* CRISPRi (Figure 2 B and C and) confirming that both knockdown methods result in decreased *miR26a* expression. The knockdown is dose-responsive; higher morpholino doses lead to a 1.8-fold reduction in *miR26a* expression (Supplemental Figure 3 A and B). In support of *smad1* being a target of *miR26a* in vivo, *miR26a* knockdown resulted in increased *smad1* mRNA expression in 2dpf and 4dpf injected embryos as compared to controls by in situ hybridization (Figure 2D). Upregulation of *smad1* was more prominent in the head of *miR26a* morphants and CRISPRi knockdown embryos, with a localization similar to where *miR26a* is expressed most strongly (Figure 1B-C). At 4dpf in both morphants and CRISPRi fish, *smad1* expression is upregulated in the pharyngeal region, with staining in the ventral aorta, aortic arches and bulbous arteriosus.

**Figure 2:**
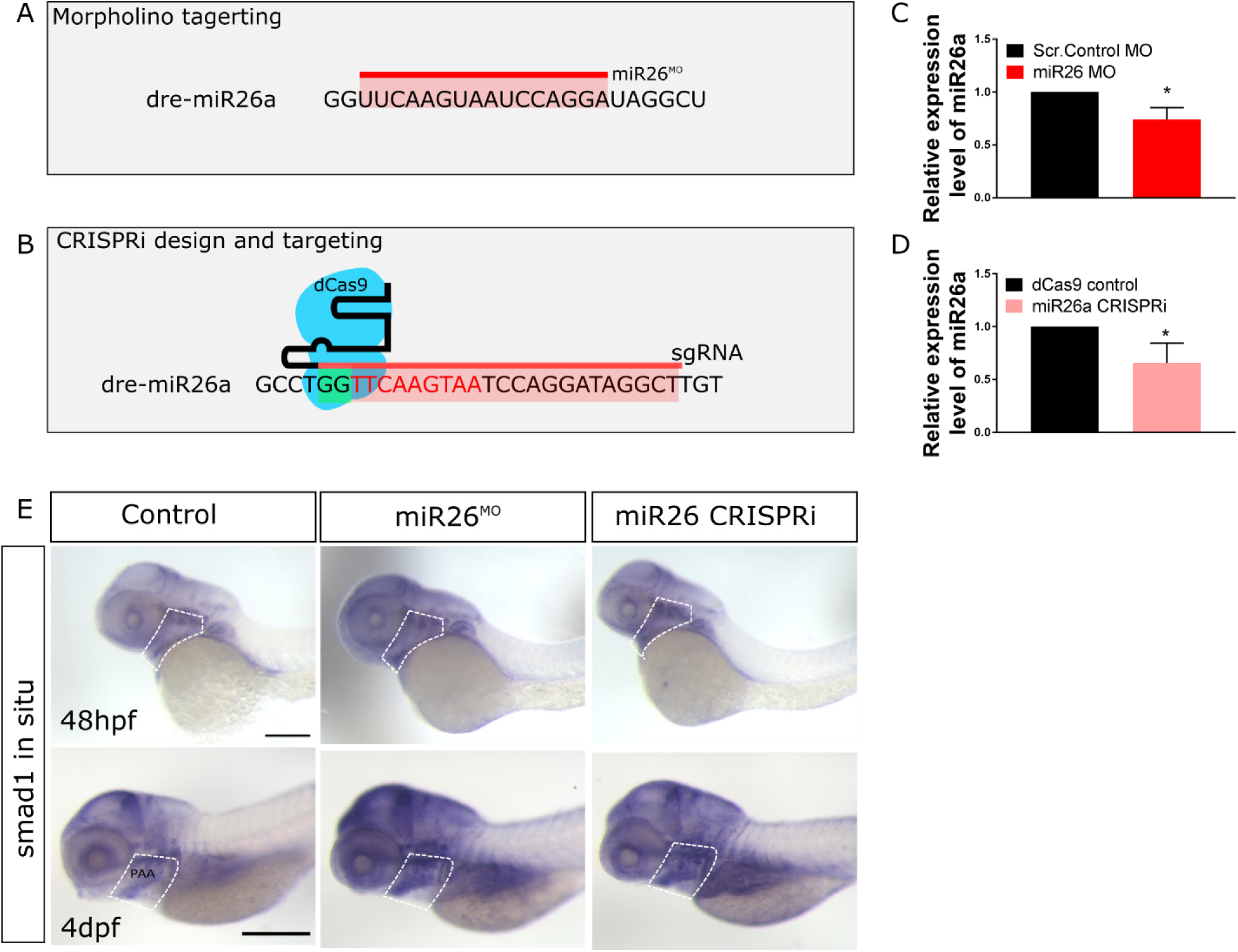
miR26 knockdown increases *smad1* expression. A and B) Schematic of miR26 transient knockdown methods. C and D) qPCR of relative expression of *miR26a* in morpholino and CRISPRi injected embryos at 2dpf, values are means of 3 replicates and normalized to miR130 expression. Error Bars = SD;*P=<0.05. E) Whole-mount in situ hybridization staining for *smad1* at 2dpf and 4 dpf shows increased expression of *Smad1* in miR26 knockdown embryos particularly in the ventral aorta, aortic arches and pharyngeal region (dotted outline). Scale bar= 500µm

**Figure 3.**
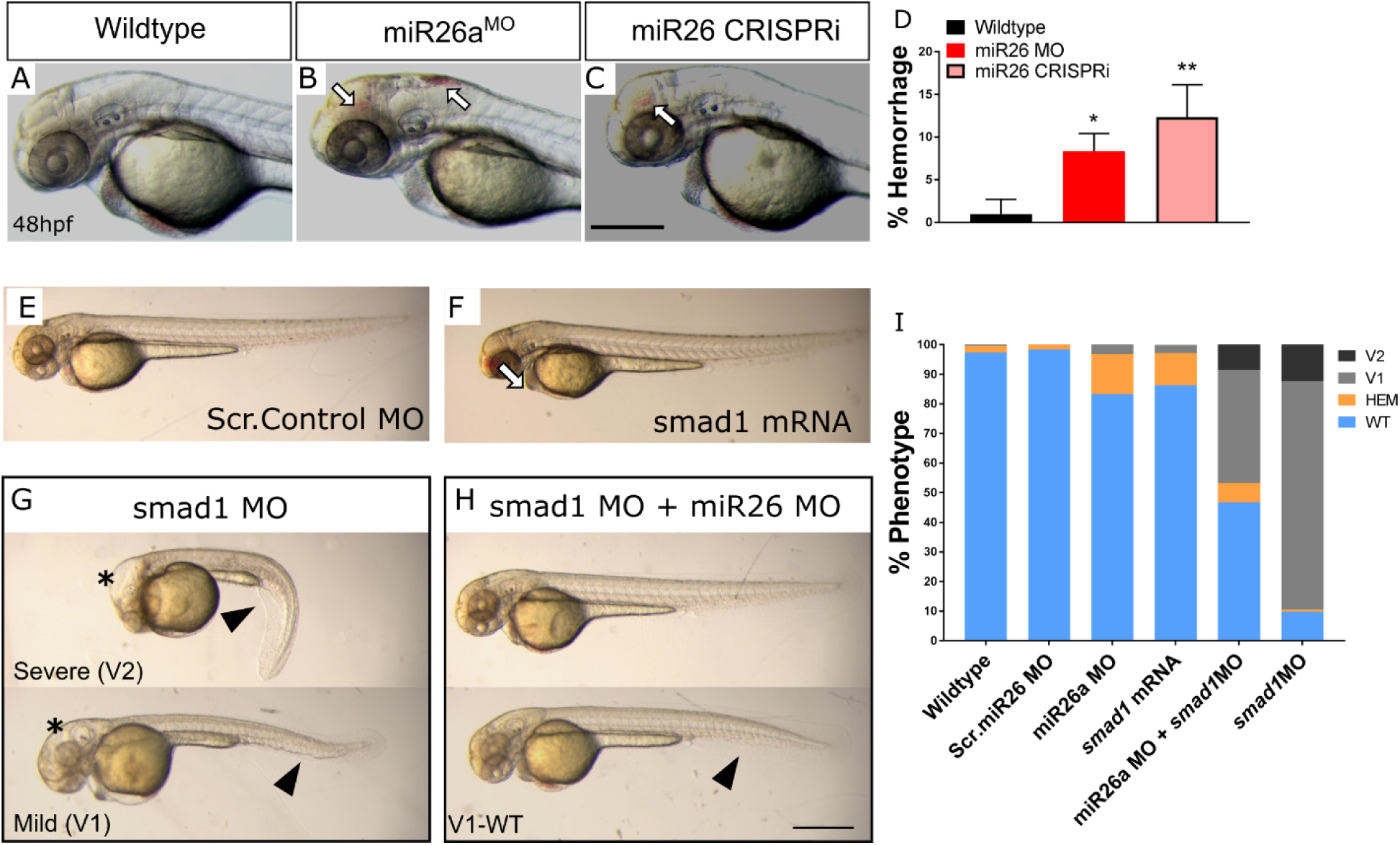
Increased levels of *smad1* result in defects in the vascular system and body axis. A-C) Representative 2dpf *miR26a* knockdown embryos with hemorrhage, as indicated by arrows. D) Quantification of average rates of hemorrhage. (Error bars = SD Unpaired t test, *miR26a* MO *p<0.01 and mi26 CRISPRi **p<0.001 as compared to WT, N=3, Wildtype =224, miR26 MO = 124, miR26 CRISPRi =180). E-F) Representative body morphology with *smad1* overexpression. G-I) *miR26a* and *smad1* double knockdown experiments. G) Representative 2dpf *smad1* MO embryos with mild (V1) and severe (V2) ventralization phenotypes. H) Representative 2dpf double *miR26a* and *smad1* knockdown embryos with rescued hemorrhage and normal body axis showing only mild (V1-WT) ventralization phenotypes. I) Quantification of observed phenotypes in single and double knockdown experiments (N=4, total n wildtype=193, Scr. Control MO = 157, *smad1* MO = 175, *smad1* mRNA = 95, miR26 MO =190, and miR26 MO + *smad1* MO = 190. Two Way ANOVA of hemorrhage phenotype. Significance: Wildtype/Scr. Control MO vs. *miR26a* MO p=0.0001 Wildtype/Scr. Control vs. *SMAD1* mRNA p=0.0001 *miR26a* MO vs. miR26 MO+ *SMAD1* MO p<0.0001 Two Way ANOVA of V1 phenotype: Wildtype/Scr. Control MO; vs. *SMAD1* MO p<0.0001 vs. miR26 MO+ *SMAD1* MO p<0.0001. Error Bars = SD. Scale bar= 500µm

In a complementary approach we injected a *miR26* mimic to overexpress *miR26a* (Supplemental Figure 2) and observed a marked increase in *miR26a* expression by qPCR and in situ hybridization (Supplemental Figure 2A, C and D). Overexpression of *miR26a* resulted in mildly dorsalized embryos by 48hpf with heart edema, dorsal axis defects and poor circulation (Supplemental Figure 2D), suggesting overexpression of *miR26* disrupts pathways such as BMP that pattern early embryonic axes.

### Increased levels of *smad1* lead to vascular stability defects

At 2dpf, *miR26a morphants* have an average 13±2% hemorrhage (Figure 3 B and D) and CRISPRi embryos have an average 15±1% hemorrhage (Figure 3 C and D). The phenotype is dose-dependent as higher doses of morpholino lead to an increase in hemorrhage to 40% (Supplemental Figure 3 C and D). As *smad1* is a known target of *miR26a*, but upregulation of *smad1* has not been previously connected to vascular stability defects, we overexpressed *smad1* by injection of mRNA into single cell stage embryos. Injection of *smad1* mRNA alone resulted in significantly higher hemorrhage rate of 22% in injected embryos as compared to uninjected controls (Figure 3F and I). Next, as *miR26a* knockdown leads to increased *smad1* levels, we predicted that reduction in *smad1* would rescue hemorrhage in *miR26* knockdown embryos. We used a double morpholino knockdown strategy using co-injection of a validated *smad1* MO with the *miR26a* morpholino, both at 1ng/embryo. Double knockdown reduced hemorrhage rates to below 5% (Figure 3H, top embryo and Figure 3I). *smad1* MO alone does not result in hemorrhage; however it did result in a range of phenotypes associated with *smad1* knockdown including dorsal-ventral axis defects and hydrocephalus as previously reported (McReynolds et al. 2007). *smad1* knockdown led to 80% of embryos with a mild (V1) ventralized defect and 15% with a more severe (V2) (Figure 3G and I). Both defects were reduced in double knockdown embryos (Figure 3 G and H, bottom embryos). Thus reducing *miR26a* levels by either MO or CRISPRi knockdown *in vivo* leads to an increase in *smad1* levels. Increasing *smad1* leads to a vascular stability defects, while reducing Smad1 levels in *miR26a morphants* normalizes the phenotype.

### Loss of *miR26a* leads to increased phosphorylated Smad1 in endothelium

We next tested whether the increased expression of *smad1* mRNA in *miR26a* knockdown embryos was also indicative of enhanced Smad1 activation by phosphorylation. Wildtype immunostaining showed pSmad1/5/9 is highest within the endothelium and not vSMCs (Supplemental Figure 1 B-B’). Using *Tg(fli:EGFP*^*y7*^*; kdrl:mCherry)* transgenic embryos we counted the number of pSmad1 positive endothelial nuclei in *miR26* morphants and *miR26* CRISPRi knockdown embryos along the ventral aorta and from the first aortic branch of the PAA to the beginning of the bulbous arteriosus (Supplemental Figure 4). We found no significant difference in the number of endothelial cells as indicated by *Fli:nEGFP*^*y7*^ positive nuclei between the injected groups (Figure 4 A-D), however there is a significant 20% increase in the proportion of pSmad1 positive:*Fliy7:nEGFP* nuclei double positive endothelial cells in *miR26* knockdown embryos as compared to controls, with an average of 60+3.1 % and 64.9+3.9 % in *miR26* morphants (Figure 4 F,J and N) and CRISPRi embryos (Figure 4 H,L and O), respectively and 42-47% in controls

**Figure 4:**
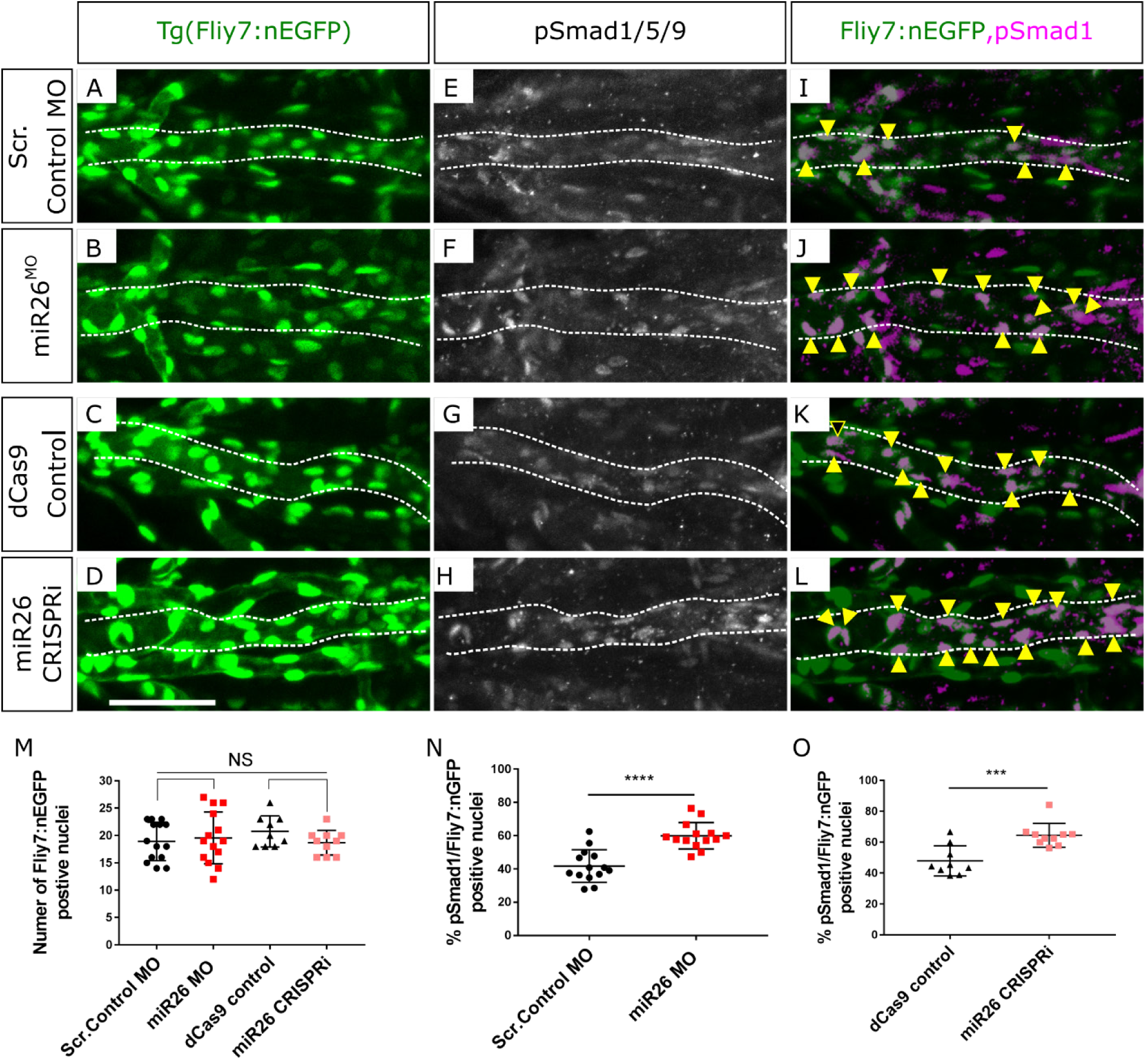
mir26 knockdown embryos have increased endothelial p*Smad1*. Ventral view confocal projections of the 4dpf ventral aorta (dotted outline). Endothelial nuclei (*fli*^*y1*^:GFP; A-D, arrowheads) and p*Smad1*/5/9 (white E-H) and overlay (magenta, I-L) in 4dpf Scr. Control (A,E,I), *miR26a* MO (B,F,J), dCas9 control (C,G,K) and *miR26a* CRISPRi (D,H,L) embryos. Yellow arrowheads indicate p*smad1* + *Fliy7*:nEGFP double positive nuclei in the ventral aorta. M) Quantification of total number of Fliy7:nEGFP nuclei in the ventral aorta. N and O) Quantification of the proportional percentage of p*smad1* positive/Fliy7:nEGFP positive nuclei in miR26 morphants and miR26 CRISPRi embryos. N=3 experiments, total embryos Scr. Control MO = 14, miR26 MO =14, dCas9 control=9 and *miR26a* CRISRPi=10. Unpaired t test, p***=0.0001 and p ****=0.00001 as compared to WT, error bars = SD, Scale bar: 50 μm.

### Loss of *miR26a* leads to increased numbers of acta2-positive vSMCs and a differentiated morphology

As both *miR26a* and *smad1* are implicated in controlling vSMC differentiation, we investigated if there was a functional consequence of the increased endothelial BMP signalling on vSMCs. We assayed the expression of a key set of marker genes that are highly expressed in differentiated vSMCs. Using in situ hybridization, we found that *miR26* morphant embryos have increased expression of *acta2* and *myh11a*, especially in the pharyngeal region where vSMCs first develop (Supplementary Figure 5 A-B and C-D). This is also a similar location to that of increased *smad1* staining (Figure 2D). Conversely by 4dpf miR26 mimic injected embryos had reduction in *acta2* and *sm22* expression (Supplementary Figure 5 E-G and H-I).

Taking advantage of live reporters, we quantitated vascular smooth muscle of the ventral aorta and pharyngeal arch arteries in *miR26* morphants (Figure 5A). This assay allows us to make a number of key observations. First, the *BRE:GFP* signal appears to be enhanced in *miR26a* morphants (Figure 5 A’, B’ and C). Although this observation is not significant it correlates with increased pSmad1 staining (Figure 4). Second, the number of *acta2:GFP* positive cells along the ventral aorta and pharyngeal arch arteries (PAA) are increased in *miR26* knockdown embryos (33.8±1.6 vs 47.22±2.2, Figure 5 A’, B’ and D). Third, the increase in *acta2* positive cell number, is accompanied by a qualitative change in cell morphology of *miR26a* knockdown embryos. In control embryos, 25% of total *acta2* positive cells have a rounded, punctate morphology, and ‘sit’ high on the vessel wall. The remaining cells having a more mature spindle, flattened morphology. In *miR26* morphants, 13% appear rounded while the majority, 86%, are spindle like cells. This data suggests that loss of *miR26a* results in increased vSMC coverage along blood vessels that vSMCs, and a shift to a more differentiated phenotype.

**Figure 5:**
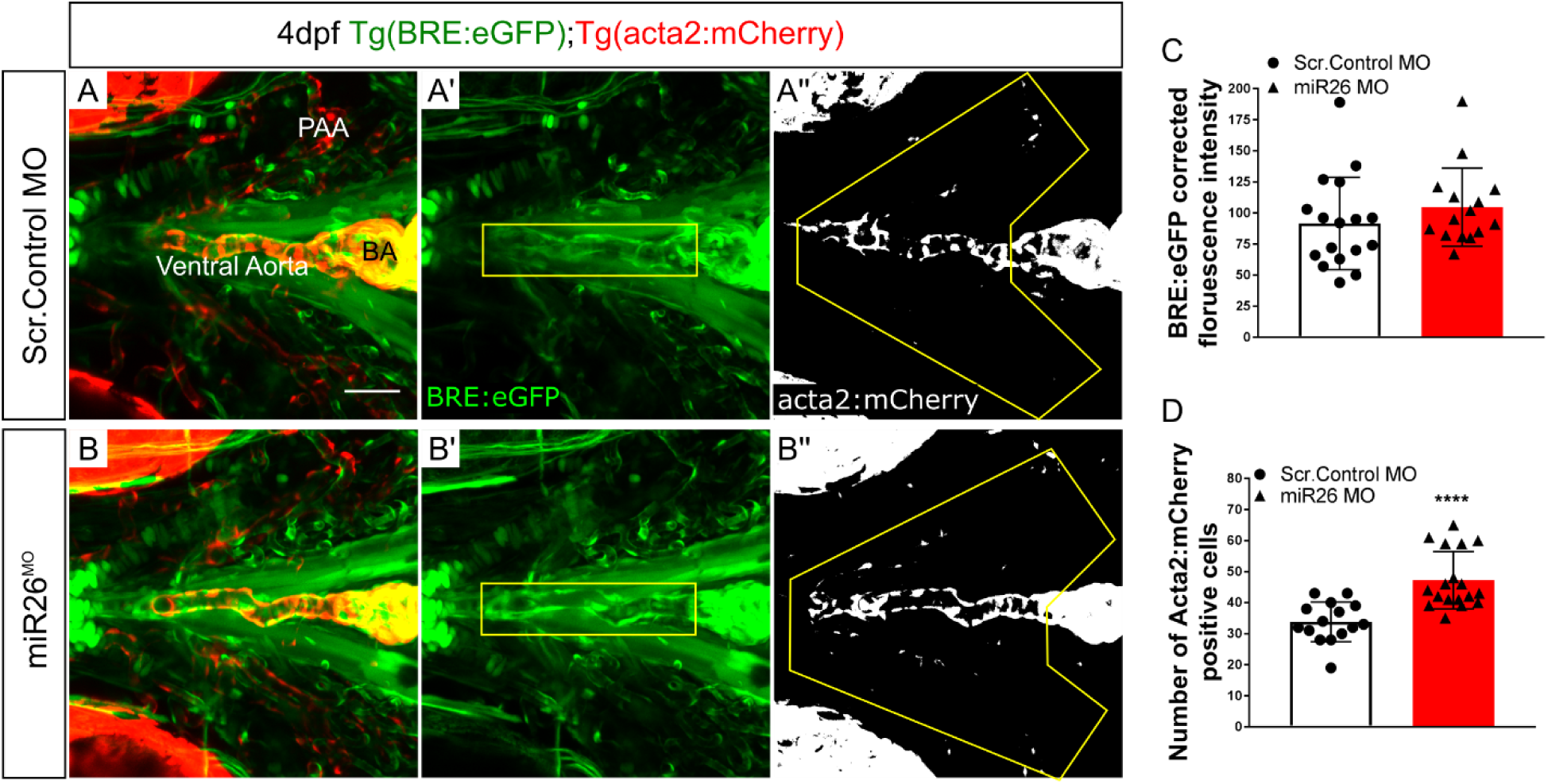
miR26 morphants have increased expression of vSMC genes and an increased number of *acta2*-positive cells. A-B) Representative orthogonal projections of ventral views of 4 dpf Tg(BRE:eGFP); Tg(*acta2*:mCherry) embryos. Wildtype embryos (A-A’’) and *miR26a* morphant embryos (B-B’’) showing qualitative upregulation of BRE:eGFP in the ventral aorta (VA) and pharyngeal arch arteries (PAA) G) Quantification of BRE:eGFP intensity along the VA, within the yellow box in B’. H) Quantification of *Acta2* positive cell number on VA and PAAs, in area outlined in B’’. Number of *acta2* positive cells is significantly increased in miR26 morphants (N=3,miR26 MO = 15, Wildtype = 13, Unpaired t test, ****p<0.0001 as compared to WT, error bars = SD).

### Inhibition of the BMP pathway reverses the effects of *miR26a* knockdown

Given that pSmad1 levels are increased within the endothelium of *miR26a* knockdown embryos and that these embryos also have an increased number of vSMC, we investigated whether blocking BMP signalling reverses the effect of loss of *miR26*a on vSMC number and differentiation. We used a pharmacological approach to reduce receptor phosphorylation of Smad1. K02288 is a selective and potent inhibitor of the BMP type I receptor activin like kinase 1 (Alk1) and Alk2. To select appropriate timepoints, the endothelium of the major blood vessels are patterned by 48hpf (Isogai et al. 2001), while vSMC coverage of the ventral aorta and PAA which begins from 52hpf (Whitesell et al. 2014). Therefore, *Tg(acta2:eGFP;kdrl:mCherry)* embryos were treated with 15µM d K02288 from 52hpf to 4dpf. Treatment at this later stage also avoids gastrulation defects and allowed us to assay the effects of BMP inhibition during a time when vSMCs first associate with blood vessels.

In wildtype embryos treatment with K02288 led to a 62% reduction in the total average number *acta2:eGFP* positive cells compared to vehicle control alone (Figure 7 A’, D’ and G. 29+1 to 11+3). In *miR26* knockdown embryos (Figure 7 B’-C’) the effects of K02288 were blunted and *acta2*-positive cell numbers were maintained at comparable levels to wildtype controls. *miR26a* morphants had a minimal 17% reduction (35+5 to 29+5) and *miR26a* CRISPRi embryos had a 12% reduction in numbers from 39+1 to 34+3 (Figure 7 E’-F’ and G). This suggests that loss of *miR26a*, and the subsequent increase in Smad1 may compensate for reduced receptor function and provide enough signal to maintain vSMC coverage. Alongside the reduction in vSMC number, we also observed an average 50% reduction in the length of ventral aorta length in K02288 treated wildtype embryos at 4dpf, from an average 159µm to 77µm (Figure 7 A, D and H). This reduction was not seen in *miR26a* morphants treated with K02288, which maintain ventral aorta length. *miR26a* CRISPRi embryos had a small, 20% decrease in length when treated with K02288 (Figure 7 B-C and E-F). Of note there was no statistical difference in ventral aorta length between *mir26a* knockdown and control embryos, which supports our finding that EC cell number is not affected. Taken together, our data indicates blocking BMP signalling effectively reduced vSMC coverage and affects endothelial development in the ventral aorta. *miR26a* knockdown animals have minimal changes in vSMC coverage in response to BMP inhibition.

**Figure 6:**
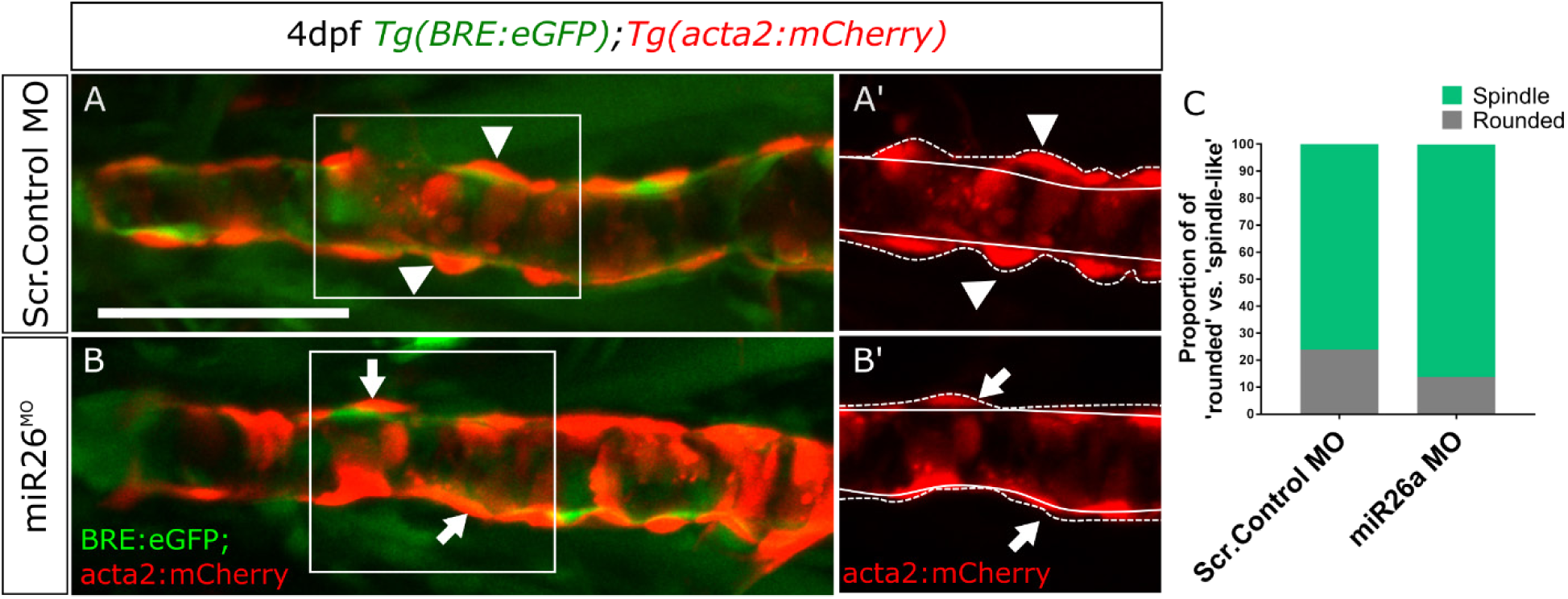
Trend to a more mature morphology of vascular smooth muscle cells in miR26 morphants. A-B) vSMC morphology in 4dpf embryos, insets A’ and B’ show punctate (arrowheads) and spindle-like (arrows) morphology. Dashed lines indicate height of cells from the endothelium. C) Quantification of proportion of punctate vs spindle shaped *acta2* positive cells in wildtype and miR26 morphants. N=3 experiments with total embryo n, miR26 MO = 15, Wildtype = 13. Scale bar= 50µm

**Figure 7:**
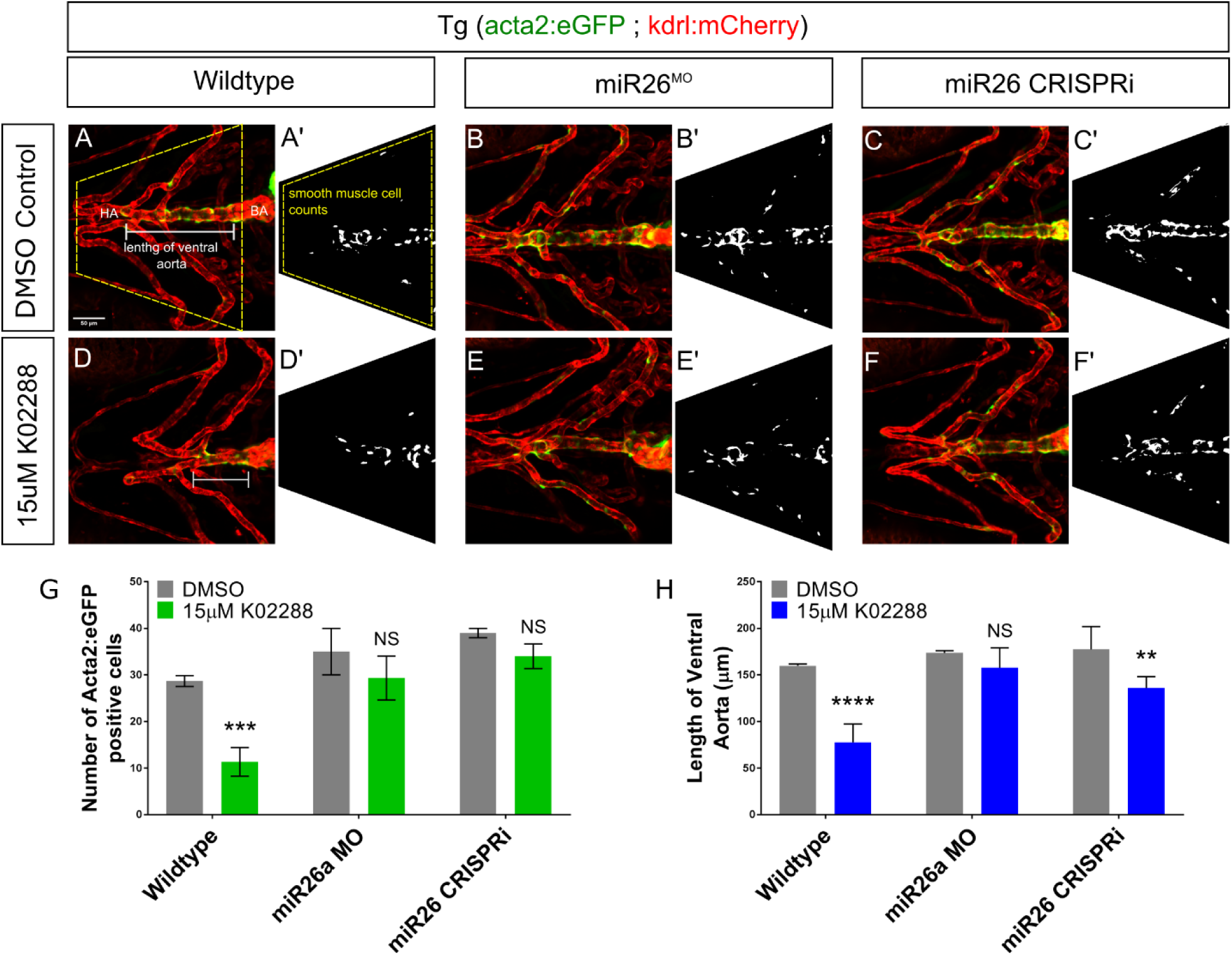
miR26 controls vSMC differentiation via *smad1*-mediated BMP signaling. Ventral aorta showing endothelial (red) and smooth muscle (green) cells in miR26^MO^ or CRISPRi-injected embryos treated with small molecule inhibitors from 52 hpf to 4 dpf. A-C) DMSO-treated control embryos D-F) 15µM K02288 treated control embryos. (A,D), miR26 morphant (B,E), miR26 CRISPRi knockdown (C,F). A’-F’) Threshold adjusted images of *acta2*-eGFP expression. (N= 3, 8-9 embryos per treatment group. One Way ANOVA, **, ***, ****p=0.001-0.00001. Scale bar= 50µm

## Discussion

Compromised structural vascular integrity, vessel weakening and rupture (hemorrhage) can result from aberrant BMP signalling (Dzau et al. 2002; Milewicz et al. 2010; Nebbioso et al. 2012). Hemorrhage ultimately results from weak endothelial junctions, however defects in mural cell coverage (attachment and ECM secretion) are implicated in the pathological progression of vascular diseases. We show that the underlying endothelium of the ventral aorta in zebrafish has activated pSmad1 at 4 dpf, but that pSmad1 is not detectable in mural cells. At a stage when mature vSMC are normally present, embryos with loss of *miR26a* have upregulation of pSmad1, increased vSMC coverage and a change in vSMC morphology, with no observable changes in the number, or morphology of the pSmad1-expressing endothelial cells. We show that inhibition of BMP signalling reduces both vSMC coverage and the length of the ventral aorta while dual *miR26* knockdown and BMP receptor inhibition leads to a rescue such that animals maintain normal vSMC number, length of the ventral aorta, and vSMC coverage. We therefore suggest that *miR26a* modulates BMP signalling in endothelial cells to control vSMC differentiation via yet unidentified paracrine mechanism’ paracrine mechanism, and that *miR26a* functions to fine tune endothelial signals to the vSMCs (Figure 8).

**Figure 8:**
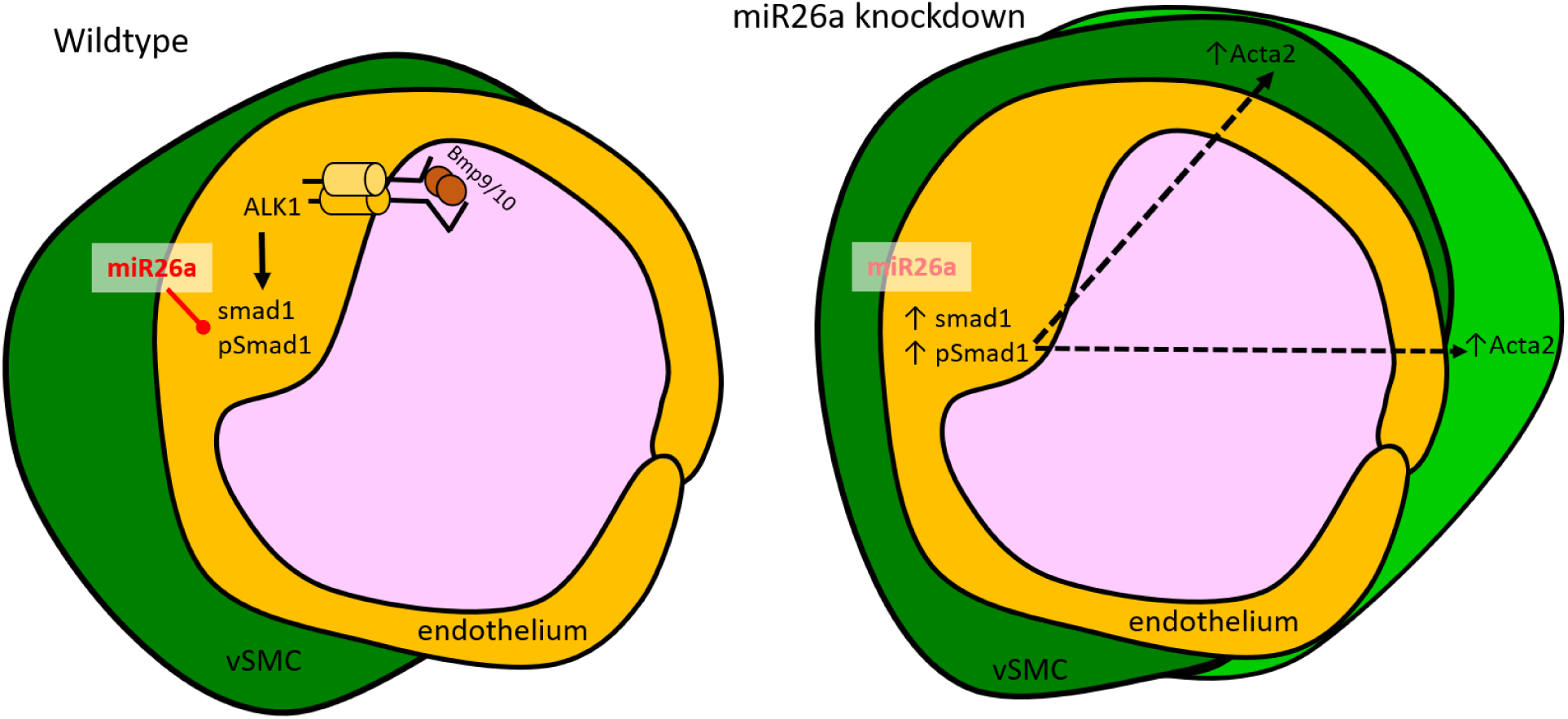
Proposed mechanism by which *miR26a* modulates BMP signaling to promote SMC phenotype via interactions with endothelial cells. miR26 modulates vascular stability by directly targeting *smad1*. At developmental stages when smooth muscle appears, the endothelium has active BMP signaling. Loss of miR26 results in increased BMP signaling in endothelial cells where *smad1* becomes phosphorylated. Increased pSmad1 in endothelial cells leads to increased *acta2* and increased *acta2*-positive vSMC cell number, while blocking BMP signaling leads to a decrease of both.

Studies in cultured vSMC have suggested that *miR26a* controls Smad1-mediated BMP signalling within SMCs to modulate their phenotype (Albinsson et al. 2010). However, these *in vitro* studies do not address whether the levels of pathway activation in vitro are relevant to tissues in vivo. Additionally, data collected from *in vitro* culture systems does not address the role of cell to cell communication (autonomous and non-autonomous signalling) that is critical *in vivo* (Gaengel et al. 2009; Owens et al. 2004). We therefore sought to use a live model of vascular development with intact tissue and cellular contexts. Using a BMP-reporter transgenic fish, we found that during normal development, and under physiological conditions, vSMC directly contact BRE-positive and pSmad1 positive endothelial cells but have undetectable BRE or pSmad1 signal in themselves. This important finding suggests that endothelial cells are key for responding to physiological BMP signals.

Treatment with K02288, a potent ALK1/2 inhibitor, significantly reduced both acta2-positive vSMC coverage and reduced the length of the ventral aorta. *miR26* knockdown embryos showed minimal reductions in both vSMC number or VA length, which suggest that the enhanced Smad1 activation in these embryos compensates for the receptor inhibition. ALK1, ALK2 and ALK3 are expressed in both endothelial and vSMCs (Benjamin et al. 1998; Lan et al. 2007; Roman et al. 2002; Beppu et al. 2000; Mishina et al. 1995), however our data suggests that although vSMC *in vitro* have the potential for autonomous BMP signalling, *in vivo*, and under physiological conditions endothelial Smad1 activation contributes to non-autonomous vSMC coverage. In zebrafish *Alk1* is highly expressed in the endothelium by 36hpf (Laux et al. 2011). *Violet beauregarde (vbg*^*ft09e*^*) alk1* loss of function zebrafish mutants develop striking cranial vessel abnormalities by 2dpf due to increased endothelial cell (EC) proliferation (Roman et al. 2002). *vbg*^*ft09e*^ are also unable to limit the diameter of arteries carrying increasing flow from the heart (Corti et al. 2011). Based on our data involving indirect control by endothelial signalling, we would predict there is a defect in vSMC recruitment when Smad1 signalling downstream of Alk1 is disrupted in these mutants, although this remains to be tested.

Our data suggests that the normal function of *miR26* is to reduce Smad1 protein activation within the endothelium, and indirectly inhibit vSMC differentiation in early development. Endothelial and mural cells signal through several paracrine pathways to stabilize vessels including the PGFRβ, angiopoietin, *TG*Fβ, and notch pathways to modulate cell-to-cell communication (Mack 2011; Winkler et al. 2011). In one of the most important pathways for mural cell recruitment, endothelial cells release PDGF-BB ligands that acts on the mural cell receptor PDGFR-β to initially recruit vSMCs to blood vessels (Benjamin et al. 1998; Lindahl et al. 1997). There is evidence that *miR26* is modulated by PDGF-BB signalling (Yang et al. 2017). Yang et al, 2017, recently showed that neointimal hyperplasia results in elevated levels of PDGF-BB and is associated with upregulation of *miR26* and the accumulation of vSMC at sites of injury, indicative of increased proliferation. Furthermore, treatment of primary mouse aortic vSMCs with *miR26a* mimic drives cells to a synthetic vSMC state (Yang et al. 2017). Currently the link between increased PDGF-BB and *miR26* is unknown, however because PDGF-BB is primarily released by endothelial cells, it should be explored whether it is activated downstream of Smad1 activation.

While we found increased differentiation of vSMCs at the later stage 4 dpf timepoint, at 48hpf loss of *miR26* results in hemorrhage. The 2dpf to 4dpf window is a common window for vascular instability phenotypes to emerge in zebrafish (Zheng et al. 2010; Liu et al. 2007; Montero-Balaguer et al. 2009; High et al. 2008). BMP signalling is initiated in endothelium at this timepoint and perturbations can affect endothelial cell junction development (Winkler et al. 2011). We have previously shown mural cells present around vessels by 2dpf, although they are mesenchymal and immature(Liu et al. 2007). These cells express PDGFRβ but have no expression of mature vSMC markers (Ando et al. 2016), suggesting the 48hpf timepoint represents a critical window for vascular mural cell attachment to endothelium and differentiation to a mature phenotype. Paracrine signals from endothelium regulated downstream of *miR26* could affect mural cell differentiation, leading to vascular instability.

As critical modulators of vascular cell function and with roles in cell differentiation, contraction, migration, proliferation and apoptosis, microRNAs are attractive targets of therapeutic treatments aimed at modulating the vSMC phenotypic switch. Specific to *TG*F-β/BMP signalling, the mir-145/143 family has direct involvement in SMC differentiation by repressing the Klf4 to induce a contractile morphology and reduced rates of proliferation (Cordes et al., 2009). miR-21 controls vSMC differentiation through cross-talk with miR-143/-145 (Sarkar et al., 2010) and by mediating *TG*F-β/BMP induction to promote miR-21 cleavage to its mature form and a more contractile phenotype (Figure 5). *miR26* is unique in this group in that it represses smooth muscle differentiation to mature states, likely via a paracrine signalling from endothelial cells. As drug delivery to the endothelium is relatively straightforward, modulation of *miR26a* might be therapeutically useful for post-transcriptional control of key genes involved in vSMC phenotypic switching.

